# Proteome Profiling of Cerebrospinal Fluid Reveals Novel Biomarker Candidates for Parkinson’s Disease

**DOI:** 10.1101/2021.07.22.453322

**Authors:** Ozge Karayel, Sebastian Virreira Winter, Shalini Padmanabhan, Yuliya I. Kuras, Duc Tung Vu, Idil Tuncali, Kalpana Merchant, Anne-Marie Wills, Clemens R. Scherzer, Matthias Mann

## Abstract

Parkinson’s disease (PD) is a growing burden worldwide, and despite ongoing efforts to find reliable biomarkers for early and differential diagnosis, prognosis and disease monitoring, there is no biofluid biomarker used in clinical routine to date. Cerebrospinal fluid (CSF) is collected often and should closely reflect structural and functional alterations in PD patients’ brains. Here we describe a scalable and sensitive mass spectrometry (MS)-based proteomics workflow for CSF proteome profiling to find specific biomarkers and identify disease-related changes in CSF protein levels in PD. From two independent cohorts consisting of more than 200 individuals, our workflow reproducibly quantified over 1,700 proteins from minimal sample amounts. Combined with machine learning, this identified a group of several proteins, including OMD, CD44, VGF, PRL, and MAN2B1 that were altered in PD patients or significantly correlate with clinical scores, indicative of disease progression. Interestingly, we uncovered signatures of enhanced neuroinflammation in patients with familial PD (LRRK2 G2019S carriers) as indicated by increased levels of CTSS, PLD4, HLA-DRA, HLA-DRB1, and HLA-DPA1. A comparison with urinary proteome changes in PD patients revealed a large overlap in protein composition PD-associated changes in these body fluids, including lysosomal factors like CTSS. Our results validate MS-based proteomics of CSF as a valuable strategy for biomarker discovery and patient stratification in a neurodegenerative disease like PD. Consistent proteomic signatures across two independent CSF cohorts and previously acquired urinary proteome profiles open up new avenues to improve our understanding of PD pathogenesis.

## INTRODUCTION

Parkinson’s disease (PD) is the second most common neurodegenerative disease, affecting millions of people worldwide and a strikingly increased incidence with age [1, 2]. The hallmark pathology of PD is well-characterized as a-synuclein aggregates and dopamine neuronal loss [2], however, molecular events that trigger PD are not fully understood. As a result, all current treatments only target symptoms, without slowing or reversing disease progression [3]. Detecting protein level alterations in PD could provide insight into the underlying mechanism of disease and aid in the development of novel therapeutics.

The majority of PD cases are idiopathic while for some a genetic linkage is apparent [3–5]. Among several gene alterations causing genetic PD, mutations in the LRRK2 gene, including the most common G2019S mutation, are the most frequent genetic cause of autosomal dominant PD [6, 7]. Importantly, all PD-associated mutations activate LRRK2 kinase activity, offering a promising therapeutic target for PD by inhibiting this function [7–9]. Better understanding of pathophysiology of LRRK2-PD and identification of related biomarkers would facilitate the development and application of LRRK2-targeted therapies. It is also essential to determine whether pathogenic mechanisms associated with LRRK2-PD may be present at the prodromal stage in non-manifesting LRRK 2 mutation carriers and in a subset of idiopathic PD.

Cellular and animal model studies indicate that LRRK2 is involved in regulation of several pathophysiologic processes, including autophagy, endolysosomal membrane/vesicular trafficking and immune responses, with at least some of the effects mediated through phosphorylation of a subgroup of Rab GTPase [10–13]. Different PD-associated genes contribute to pathogenesis through common molecular pathways, but additional pathological changes in multiple pathways also occur [14, 15]. While cell biology research on molecular pathways affected by PD-associated genes is critical for insights into disease mechanisms, a complementary approach would be to examine molecular changes in bio specimens from deeply phenotyped LRRK2 and idiopathic PD cohorts compared to controls. We thus reasoned that an unbiased, mass spectrometry (MS)-based proteomics analysis of cerebrospinal fluid (CSF) from heathy controls, PD individuals with and without LRRK2 G2019S mutation and non-manifesting LRRK2 G2019S carriers would be a promising avenue to uncover much-needed biomarkers. Compared to other biofluids, CSF has major advantages in the study of neurodegenerative and neuroinflammatory diseases as it surrounds the brain and the spinal cord and contains brain-derived proteins and other biomolecules [16, 17]. Therefore, it likely reflects disease-related pathology of the brain and spinal cord more accurately, providing important and novel information.

Mass spectrometry (MS)-based proteomics has become a very powerful technology for unbiased detection of differences in protein abundance levels in healthy individuals and patients, and thus – in principle - an ideal tool for biomarker discovery . However, proteomic analysis of body fluids including CSF has been challenging due to low protein concentration in CSF combined with the high dynamic range of protein abundances, generally resulting in low quantification precision, throughput and limited proteome depth [18, 19]. In particular, the presence of highly abundant proteins in body fluid specimens such as albumin substantially interferes with reproducible quantification of less abundant proteins, resulting in missing values across samples [20].

Recent advances in the proteomics field, from automated sample preparation to new more sensitive MS instrumentation and data acquisition methods such as data-independent acquisition (DIA) and processing software, allow substantial proteome coverage and precise quantitation in a single run [21–25]. Our group has combined these advances to develop a streamlined and highly reproducible workflow resulting in a large number of consistently measured and biologically meaningful proteome changes in various biofluid/tissue specimen in a variety of clinical cohorts [18, 19, 26–29].

In this study, we extended our optimized pipeline of biomarker discovery to analyze more than 200 CSF samples from two independent cohorts in which we detected on average over 1,700 proteins from minimal CSF sample amounts. By employing co-variate (ANCOVA) analysis and machine learning, we identified unique protein signatures whose abundance was changed in PD patients compared to healthy controls. We also identified several proteins that were specifically upregulated in LRRK2 G2019S carriers. Our study demonstrates that modern MS-based proteomics is a powerful technology for biomarker discovery in bio fluids. Our analysis provides potential biomarkers of PD as well as insights into biological pathways associated with PD and/or LRRK2 mutations.

## RESULTS

### Overview of PD cohorts for CSF proteome profiling

To investigate how PD affects the CSF of patients and to identify potential biomarkers, we employed the ‘rectangular’ biomarker discovery strategy which aims to discover discriminating proteome signatures using rather large sample sizes in both discovery and validation cohorts [18, 19]. Applying this approach, we analyzed CSF samples collected from 215 individuals from two independent cohorts including 113 healthy controls (HC) and 102 PD. The first cohort consisted of 94 CSF samples from the Harvard Biomarkers Study (hereinafter referred to as ‘HBS cohort’) [30–33] . The second cohort was a subset of biobanked CSF samples from the Michael J. Fox Foundation for Parkinson’s Research (MJFF)-funded *LRRK2* Cohort Consortium (LCC) (hereinafter referred to as ‘LCC cohort’). More information about the cohorts are summarized in **Table 1** and **Figure S1**.

**Table 1.** Demographics of all participants

| **LCC cohort** | LRRK2-/PD-<br>(n = 31) | LRRK2+/PD-<br>(n = 35) | LRRK2-/PD+<br>(n = 34) | LRRK2+/PD+<br>(n = 21) |
| --- | --- | --- | --- | --- |
| Sex (male/female) | 13/18 | 19/16 | 23/11 | 10/11 |
| Age at collection, mean (SD) | 54.2 (13.9) | 52.1 (15.2) | 57.2 (11.2) | 62.2 (10.8) |
| Age at onset, mean (SD) | n/a | n/a | 51.1 (11.4) | 53.9 (11.4) |
| UPDRS III | n/a | n/a | 25.9 (9.4) | 23.1 (15.6) |
| MOCA score | 27.3 (2) | 26.2 (2.5) | 26.5 (3) | 24.7 (3.9) |

| **HBS cohort** | PD- (n = 47) | PD+ (n = 47) |
| --- | --- | --- |
| Sex (male/female) | 29/18 | 34/13 |
| Age at collection, mean (SD) | 56.4 (11.3) | 62.8 (8.9) |
| Age at onset, mean (SD) | n/a | 58.4 (8.4) |
| GBA (mut/WT) | 0/47 | 6/41 |
| LRRK2 (mut/WT) | 0/47 | 1/46 |
| UPDRS total | n/a | 37.5 (14.3) |
| UPDRS III | n/a | 22.1 (14.1) |
| MMSE score | 28.4 (1.8) | 28.3 (1.6) |

### Proteomic characterization of CSF samples

We have developed a robust, automated and high-throughput mass spectrometry (MS)-based proteomics workflow using a data-independent acquisition (DIA) strategy to perform proteome profiling of minimal amounts of CSF [23, 26, 34] (**Figure 1A**). This strategy has previously resulted in an unprecedented depth at high data completeness in CSF and revealed biologically meaningful proteome changes across multiple independent Alzheimer disease cohorts [26]. Here we applied our workflow to discover proteome changes in the CSF of PD patients with or without the disease-associated G2019S mutation in the *LRRK2* gene. To maximize proteome depth and coverage, we generated cohort-specific hybrid spectral libraries by merging three sub-libraries: (1) a library constructed by data-dependent acquisition (DDA) consisting of 24 fractions of pooled CSF samples; (2) another DDA library consisting of 8 fractions of extracellular vesicles enriched from pooled CSF samples; and (3) a direct-DIA library generated from the DIA analysis of all analyzed samples (meaning the library is constructed on the fly on the samples, see Methods). Matching to cohort-specific hybrid libraries of more than 5,418 and 3,167 proteins yielded 1,493 and 1,626 protein in total for the HBS and LCC cohorts, respectively, with more than 1,300 in common (**Figure 1B)**. On average, we quantified 1,357 (HBS) and 1,481 (LCC) protein groups per neat CSF sample in single runs (**Figure 1C-D, Supplementary Table 2**) and the quantified protein intensities spanned over four orders of magnitude (**Figure 1E-F**). The top ten most abundant proteins alone contributed around 38% to the total CSF proteome signals, illustrating the analytical challenges (**Figure 1E-F**).

**Figure 1.**
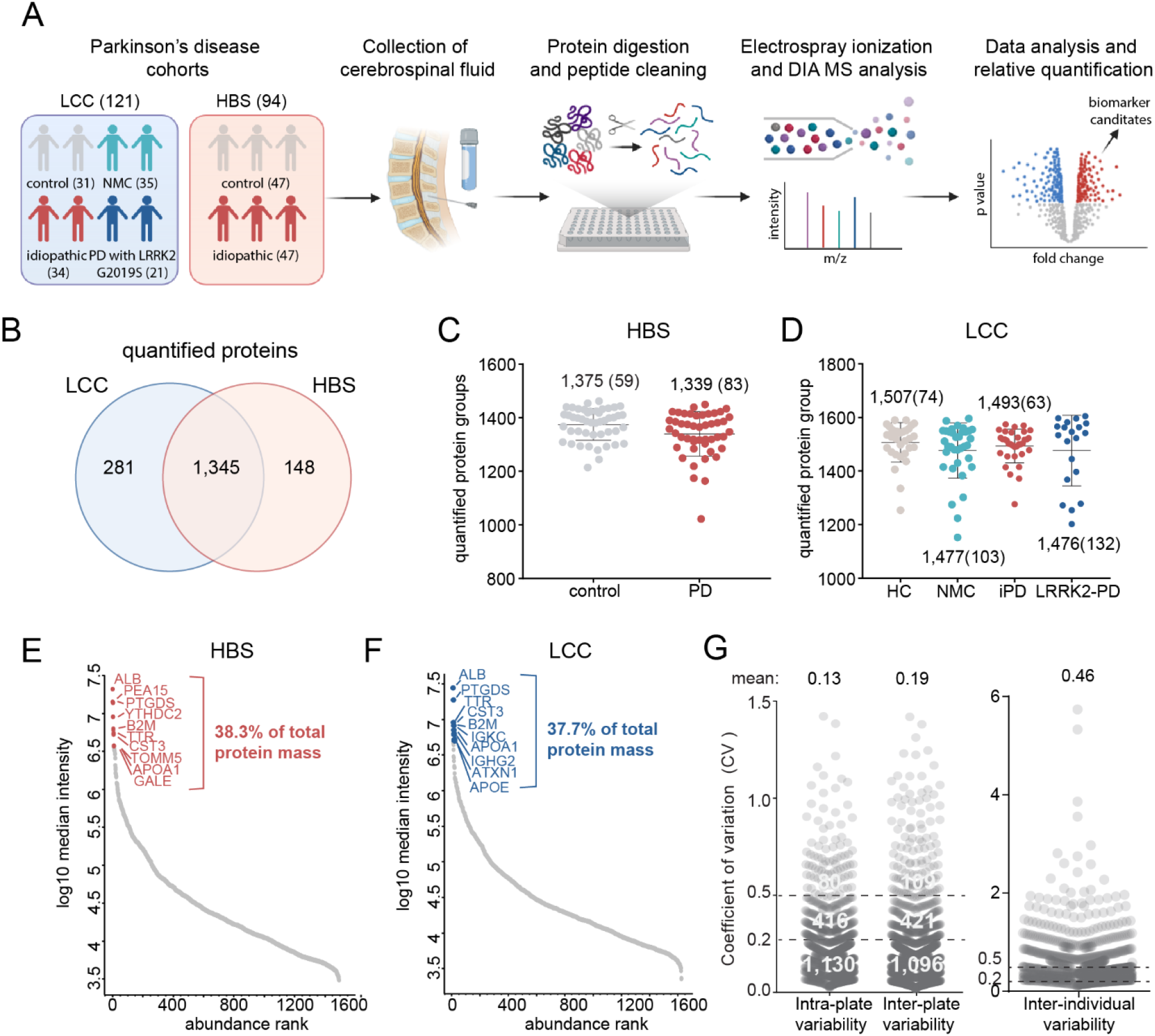
MS-based proteomic analysis of two independent CSF PD cohorts. **A**. Overview of the CSF proteomic workflow. The two cohorts comprised of either HC and iPD subject groups or HC, iPD, NMC and LRRK2 PD subject groups. CSF samples were prepared in 96-well plates using an automated liquid handling system and analyzed by LC-MS/MS using data-independent acquisition (DIA). The total number of subjects per cohort group is shown. **B.** A total of 1,345 proteins were consistently quantified in both cohorts. **C-D**. Number of proteins identified and quantified with a 1% false discovery rate (FDR) in each sample in the HBS (C) and LCC (D) cohorts. Numbers indicate mean and (standard deviation). **E-F.** Proteins identified in the HBS (E) and LCC (F) cohorts were ranked according to their MS signals, which covered about four orders of magnitude. The top ten most abundant proteins are labeled and their relative contribution to the total protein amount is indicated. **G.** Quantification precision assessed by calculating the intra- and inter-plate (between repeated measurements of the same sample) and inter-individual coefficients of variation (CVs) of all proteins. Number of proteins with a CV below and above 20% and 50% and mean CV values are shown.

We further classified the identified proteins in CSF based on their human protein atlas (HPA) annotation [35] as secreted from the cell, intra-cellular or in the cellular membranes (note that many proteins are assigned to several compartments). In total, 94% of identified proteins carried at least one annotation, of which 36% were secreted proteins while 70% and 36% were intracellular and membrane-spanning proteins, respectively (**Figure S2A**). Furthermore, the majority of quantified proteins were annotated to be enriched in brain and liver, in line with the fact that CSF surrounds the brain but is derived from blood plasma, which contains many proteins synthesized in the liver (**Figure S2B**). In summary, using minimal CSF volume, we obtained an unprecedented CSF proteome coverage for single-run analysis, thus providing a promising basis for the discovery of biomarkers in PD.

### Assessment of quantification precision and sample quality

To assess quantification precision in our study, we investigated intra- and inter-assay variabilities of our automated workflow by repeated measurements of a pooled CSF sample (**Figure S2C**), which revealed high reproducibility with around 1,100 and 1,500 proteins having inter- and intra-assay CVs below 20 and 50%, respectively (**Figures 1G and S2C-D**). The inter-individual variability is much larger with only 10% of all proteins having a CV below 20% (**Figure 1G)**. This demonstrates that the technical variability of our assay is much smaller than the biological variability - an important pre-condition for the successful discovery of novel PD proteome signatures and potential biomarkers in CSF.

Furthermore, inconsistent sample collection and handling may result in systematic bias and hamper the discovery of true biomarkers. To ensure that the quantified changes in CSF proteins are due to disease-related pathological alterations, we assessed the quality of all samples according to previously established quality marker panels for coagulation-related proteins, platelets and erythrocytes to identify samples with potential issues in pre-analytical processing [18, 19, 36] (**Figure 2A-B**). This revealed high sample-to-sample variability for the degree of contamination with erythroid-specific marker panel, presumably related to puncture related contamination with blood (**Figure 2A-B**). We flagged 14 samples (8 PD, 6 HC) from the HBS cohort and 17 samples (4 PD, 13 HC) from the LCC cohort for erythrocyte contamination and removed them from further analysis (**Figure 2C-D**).

**Figure 2.**
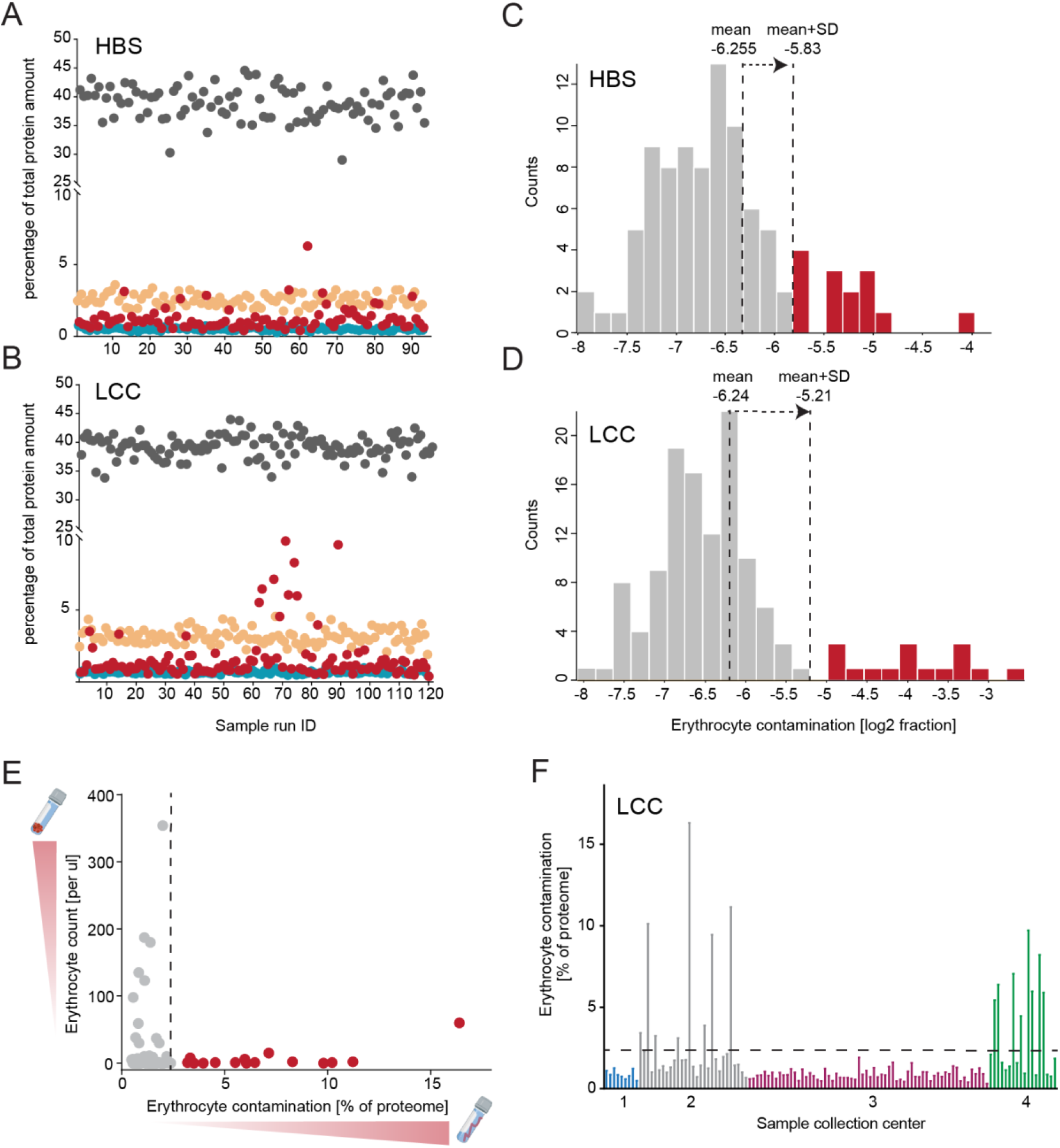
Quality assessment of two independent CSF PD cohorts. **A-B**. Assessment of study quality by determining the percentage of the summed intensity of the proteins in the respective quality marker panel and the summed intensity of all proteins in HBS (A) and LCC (B) cohorts. Erythrocyte-specific protein panel (red), platelet marker panel (turquois), coagulation marker panel (orange) and the top ten most abundant protein panel (dark grey) are included in these analyses. The proteins in each quality marker panel are listed in Supplementary Table 3. Coefficients of sample-to-sample variation for each marker panel are shown on the right. **C-D**. Histograms of log2 transformed ratios of the summed intensity of the erythrocyte-specific proteins and the summed intensity of all proteins in HBS (C) and LCC (D) cohorts. A sample was flagged for potential contamination and removed from further analysis if the ratio differed more than a standard deviation from the mean of all samples within the cohort. **E**. Comparison of erythrocyte counts in CSF following sample collection and degree of erythrocyte contamination as determined by MS-based proteomics of all LCC cohort samples. Samples colored in red were excluded from further analysis based on the distribution shown in D. **F**. Grouping samples in the LCC cohort for four sample collection centers demonstrated a high degree of contamination with erythroid-specific proteins for study centers 2 and 4, whereas there was no indication of this in study centers 1 and 3.

We found no correlation between erythrocyte counts and the amount of erythrocyte-specific proteins in the CSF (**Figure 2E**). This could be because the first draw of the CSF collection is frequently used for cell count determination, while later collected draws are stored for further analyses including proteomics. In more detail, CSF samples were centrifuged following collection to remove cells and debris, which could also explain why initially high erythrocyte counts did not affect the proteome measurements. Hemolysis occurring during sample collection, however, would only be visible in the proteome but not in erythrocyte counts. Sorting the samples by study center they were collected at, revealed a systematic bias in the sample taking and processing procedure and identified two study centers with high pre-analytical variation and a corresponding high degree of proteome contamination with erythroid-specific proteins. This further emphasizes the importance of standardized sample collection and processing procedures to minimize or avoid such biases and the advantage of unbiased proteomics to flag and remove such problematic cases.

Additionally, we further excluded a single sample from the HBS cohort that clustered far away from all other samples in a principal component analysis (PCA) and a single sample from the LCC cohort that was an outlier in proteome depth. Our subsequent bioinformatic analysis was based on the remaining 79 samples from the HBS cohort and 103 subjects in the LCC cohort to determine the impact of disease manifestation and LRRK 2 mutation status on the CSF proteome (**Figure S2E-F**).

### PD-related proteome alterations in CSF and machine learning-based classification of PD patients and controls

As PD primarily manifests in the central nervous system, its pathology should be reflected most accurately in the CSF proteome. To investigate alterations in the CSF proteome of PD patients compared to controls, we performed an ANCOVA analysis, considering age, sex and *LRRK 2* mutation status as confounding factors. Since the LCC cohort was from a multi-center study, we also included the study center as a confounding factor for this cohort. For the HBS cohort, we further included the *GBA* mutation status. Using a 5% false-discovery rate (FDR), we identified 3 significantly regulated proteins in the CSF of PD patients compared to controls in the HBS and 1 in the LCC cohort. (**Figure 3A and Supplementary Table 4**). The significantly changed proteins included CPM, OMD, RFNG in the HBS cohort and PRCP in the LCC cohort. Furthermore, osteomodulin (gene name: OMD) and the cell surface marker CD44 reached a considerable statistical specificity in both cohorts (OMD: q values of 0.063 in LCC and 0.039 in HBS and CD44: q values of 0.08 in LCC and 0.12 in HBS) (**Figure 3A)**. One reason for the small overlap between the cohorts could be a less stringent and inconsistent sample collection protocol in the LCC cohort, which itself was collected at multiple sites. Yet, the osteomodulin and CD44 proteins were robustly quantified in both cohorts and detected with three (CD44) and seven (OMD) peptides in all samples in the LCC and HBS cohorts (**Supplementary Table 4).** The levels of both proteins were significantly elevated in PD patients with mean fold changes of 1.22 (OMD) and 1.12 (CD44) in the HBS and 1.17 (OMD) and 1.07 (CD44) in the LCC cohort (**Figure 3B-E**). These results demonstrate that the rectangular strategy was able to distinguish PD-related alterations comprising a few proteins differentially present in PD patients compared to controls from cohort-specific effects in the quantified CSF proteome even in cohorts constrained by biases such as less stringent and inconsistent sample collection.

**Figure 3.**
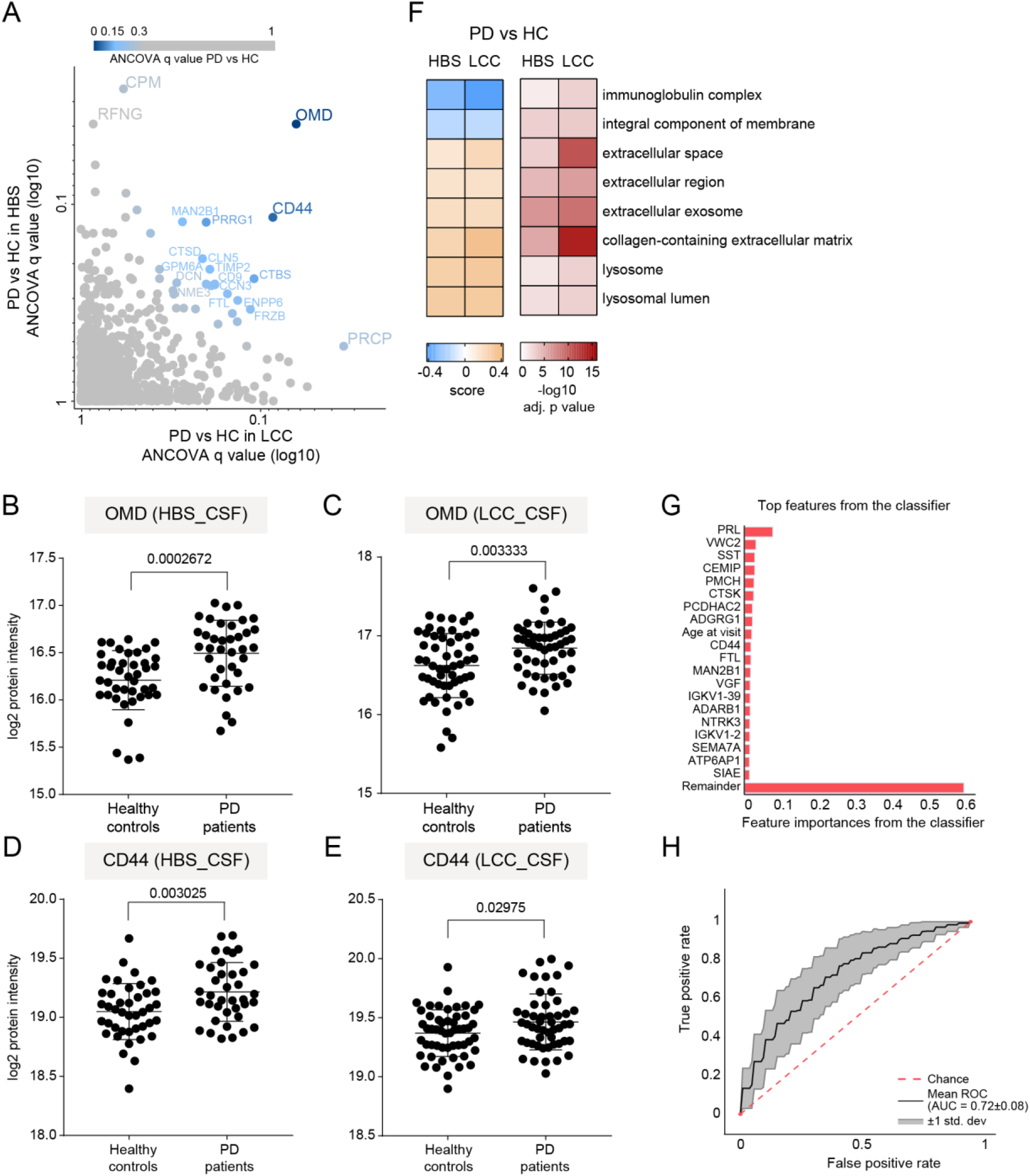
PD-related alteration in CSF proteome and machine learning-based classification of PD status. **A.** Correlation of ANCOVA q values of all proteins quantified in the CSF of PD patients compared to controls in the HBS and LCC cohorts. Color gradient is based on the mean of ANCOVA q values (PD vs HC) obtained in the LCC and HBS cohorts. **B-C.** Osteomodulin (OMD) protein intensity (log2) distribution in controls and PD patients of the HBS (B) and LCC (C) cohorts. We applied an unpaired t-test and the resulting p value is shown. **D-E.** CD44 protein intensity (log2) distribution in controls and PD patients of the HBS (D) and LCC (E) cohorts. P value from unpaired t-test is shown. **F.** Annotation enrichment of GO terms using the PD vs. HC fold-changes (5% FDR). All significantly enriched GO terms that were common in both cohorts are displayed. Terms with positive enrichment scores are enriched in PD over HC and vice versa. **G.** Top20 most important features were selected using an ExtraTrees classifier to distinguish PD+ vs. PD-individuals. **H.** Receiver operating characteristic (ROC) curve and corresponding area under the curve (AUC) statistics in 5-fold cross-validation repeated for 10 times using the XGBoost-based model to classify PD vs. HC based on protein panel in G. Random performance is indicated by the dotted diagonal red line for comparison. The gray area represents the standard deviation from the mean ROC curve. The blue lines show the values for a total of repeats with five stratified train-test splits.

To examine if PD affects particular cellular compartments and biological networks in CSF, we performed a Gene Ontology (GO) annotation enrichment analysis using the mean fold-changes of PD vs. HC [37]. The proteins elevated in the PD samples compared to the controls were enriched for categories related to lysosomal terms, further supporting the emerging role of lysosomes and mounting evidence for lysosomal dysregulation and associated a-synuclein aggregation in PD [13, 38, 39] (**Figure 3F**).

Motivated by the presence - but small number - of commonly altered proteins in both cohorts, we tested how well machine learning (ML) could discriminate PD patients from controls using the recently introduced open-source tool OmicLearn [40]. For ML purposes the number of subjects is not large which prompted us to perform a combined analysis to identify reliable signatures using machine learning. Using the ExtraTrees package, we selected the 20 most discriminating proteins for training the model (**Figure 3G**). Interestingly, among them, prolactin (gene name: PRL) was the most important feature. Although prolactin was not significantly regulated in either cohort, its release is known to be suppressed in PD patients taking dopaminergic medications [41], proving validation of our biomarker panel and pipeline. PRL did not correlate with levodopa equivalent daily doses (LEDD), provided to all PD subjects in the HBS cohorts (**Figure S3E**). Further promising candidates to classify PD vs controls were CD44, VGF and - in agreement with the lysosomal pathway enrichment we observed based on protein fold-changes of PD vs. HC-lysosomal proteins cathepsin K (CTSK) and MAN2B1. When using these features to train XGBoost, an ensemble tree-based model, with our cross-validation scheme, the mean AUC of the ROC curve was 0.72± 0.08 with sensitivity and specificity of 67 and 66%, respectively (**Figure 3H**). Taken together, our CSF proteome data, when combined with machine learning algorithms, was able to classify disease status and more importantly, identified several promising novel and PD-associated proteins that open up interesting leads for future studies.

### Impact of the pathogenic LRRK2 G2019S mutation on the CSF proteome

Given the substantial number of subjects carrying the *LRRK2* G2019S mutation in the LCC cohort, we explored whether the CSF proteome is altered by LRRK2 mutation status. We again applied an ANCOVA analysis with sex, age at sample collection, study center and PD status as confounding factors and compared the proteomes between G2019S and wild-type allele carriers. At an FDR of 5%, the mutation significantly altered the abundance of the proteins HLA-DRA, HLA-DRB 1, HLA-DPA1, CTSS, PLD4, TKT, ITGB2, PRDX3, ITIH5, CNDP1 and FAH (**Figure 4A**). Based on the peptide sequences identified in the samples, we could distinguish two forms of the protein HLA - DRB1 which were both significantly enriched in LRRK2 G0219S carriers. Furthermore, a Student’s t-test with an FDR of 5% visually confirmed the upregulation of HLA-DRA, HLA-DRB1 and HLA-DPA1 in the mutation carriers (**Figure 4A-B**). These HLA proteins as well as PLD4 and Cathepsin S (gene name: CTSS) were robustly quantified with at least 4 peptides each in the LCC cohort and their levels were significantly elevated in PD patients (>1.7-fold) (**Figure 4B-D and Supplementary Table 4**). Using this approach globally, we observed that proteins elevated in the CSF proteomes of LRRK2 G2019S carriers compared to wildtype allele carriers were enriched for categories related to immune and inflammatory responses, further supporting the substantial evidence of a close association between enhanced inflammatory response and PD [42, 43] (**Figure 4E**).

**Figure 4.**
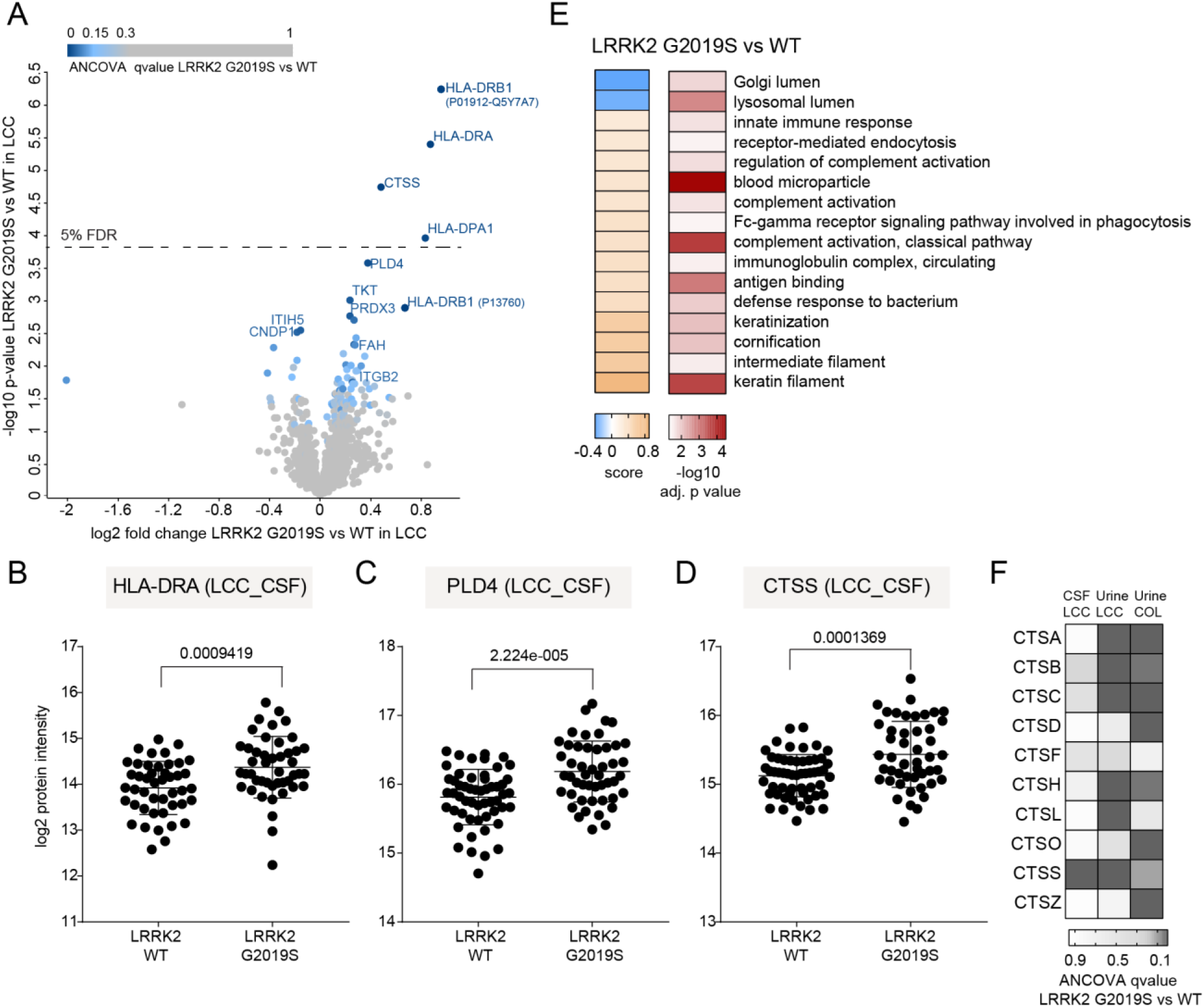
The effect of the pathogenic LRRK2 G2019S mutation on the CSF proteome. **A.** Volcano plot comparing the CSF proteomes of LRRK2 G2019S versus WT carriers. The fold-change in protein levels is depicted on the x-axis and the -log10 t-test p-value on the y-axis. Color scale is based on ANCOVA q values of the proteins differentially present in the CSF of LRRK2 G2019S carriers compared to the LRRK2 WT controls in the LCC cohort. Proteins with ANCOVA q values < 5% are labeled. **B-D.** HLA-DRA (B), PLD4 (C) and CTSS (D) protein intensity (log2) distributions in LRRK2 G2019S and WT carriers of the LCC cohort. P values of an unpaired t test are shown. **E.** Annotation enrichment of GO terms using the LRRK2 G2019S vs. LRRK2 WT fold-changes (5% FDR). Terms with positive enrichment scores are enriched in the G2019S mutation over the WT and vice versa. **F.** Heat map of ANCOVA q values of all cathepsin proteins which were quantified in both CSF and urine of LRRK2 G2019S carriers compared to the LRRK2 WT controls.

The cathepsin protein family members are proteases mediating protein degradation and turnover in endolysosomal compartments and several of them have been implicated in inflammatory diseases, lysosomal storage and neurodegenerative disorders such as PD [44, 45]. We have recently shown that the levels of multiple members of the cathepsin family including cathepsins A, B, C, D, H, L, O, S, and Z significantly increased in the urine of LRRK2 G2019S carriers in two independent cohorts [27]. These cathepsins were also detected in CSF but – in contrast to urine - only cathepsin S (CTSS) was significantly affected by the mutation status (**Figure 4F)**.

### Integration of CSF and urinary proteome profiles

We previously analyzed 235 urine samples from two independent cross-sectional cohorts including two types of controls, healthy individuals and LRRK2 G2019S carriers not manifesting the disease, and PD patients with and without the LRRK2 G2019S mutation, quantifying 2,365 urinary proteins in total [27]. Encouraged by the great depth of the acquired urine and CSF proteomes in our previous and the present study, we decided to integrate both proteomes to determine co-regulated proteins. Although we analyzed both urine and CSF subsets from subsets of the LCC cohort, there were no matching urine and CSF samples from the same individuals. More than 1,000 proteins overlapped, corresponding to 36% of all identified proteins in both biofluids (**Figure 5A**). Matching the common protein abundances revealed a clear correlation between the two biofluids (Pearson *r* = 0.49, **Figure 5B**). The most abundant proteins present in both included ALB, PTGDS, ORM1, SERPINA1, B2M and several apolipoproteins and immunoglobulins (**Figure 5B,** labeled proteins at the top right). Moreover, a Fischer’s exact test on the GO terms associated with the proteins specifically found in either urine or CSF or their common overlap revealed terms significantly enriched compared to all proteins identified in the two body fluids (**Figure 5C**). As expected, the terms related to nervous system including ‘postsynaptic membrane’, ‘memory’, ‘neuropeptide signaling pathway’, and ‘nervous system development’ were enriched among proteins exclusively present in CSF. In contrast, terms related to endosome-lysosome pathway as well as Rab protein signal transduction were enriched among urine-specific proteins, indicating that proteins of the Rab-LRRK2-pathway are highly abundant in urine, in line with our previous finding that the LRRK2 G2019S mutation strongly affects the urinary proteome [27].

**Figure 5.**
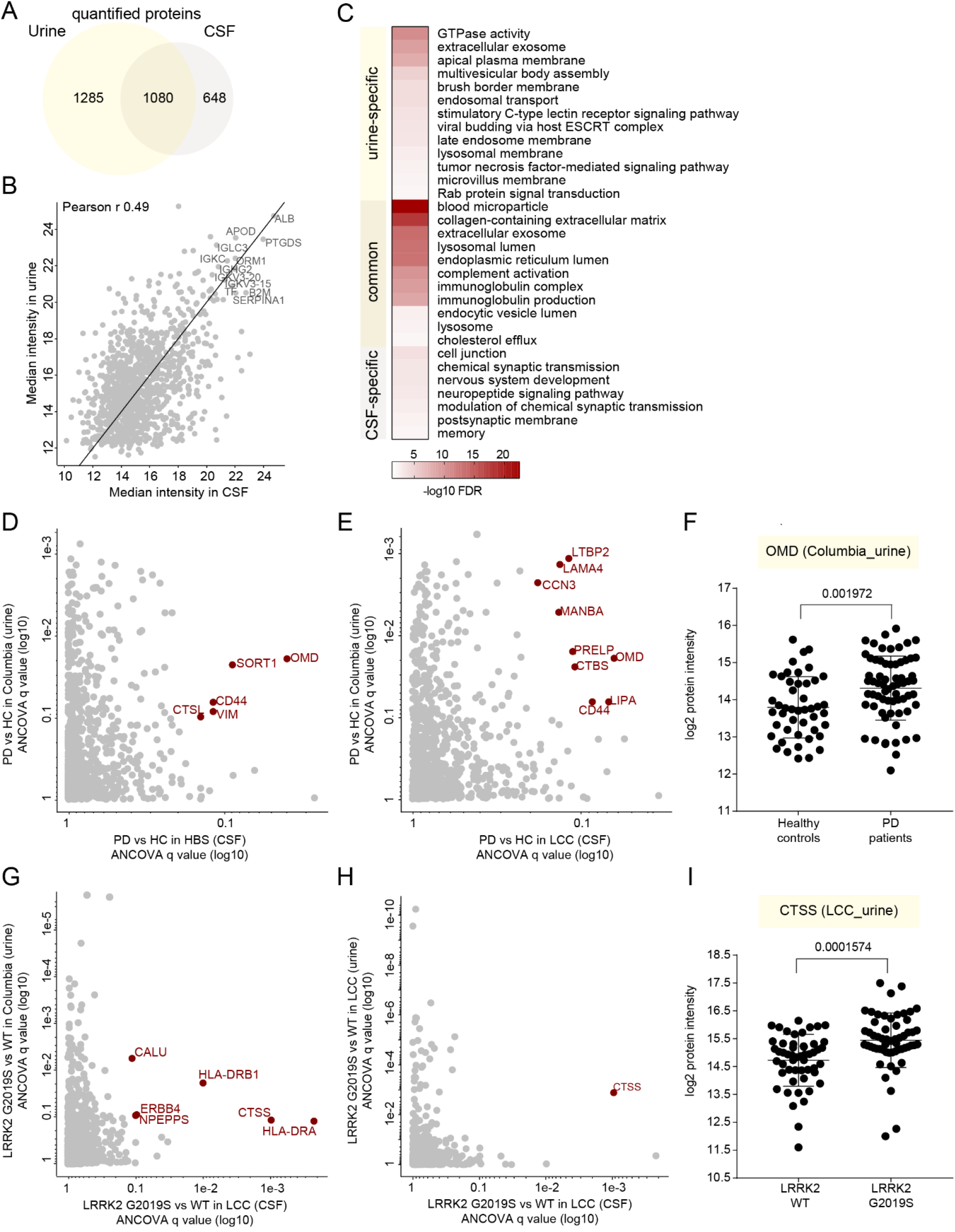
Integration of CSF and urine proteome profiles. **A.** Overlapping proteins between the CSF and urinary proteomes. **B.** CSF-urine proteome abundance map based on median MS intensities of common proteins. Highly abundant proteins in both datasets are labeled as examples. **C.** Fisher exact test to identify significantly enriched GO terms among the common and urine - and CSF-specific protein groups. Significant and non-redundant GO-terms are displayed (FDR < 5%). **D.** Correlation of ANCOVA q values of the proteins differentially present in the CSF (HBS) and urine (Columbia) of PD patients compared to controls. **E.** Correlation of ANCOVA q values of the proteins differentially present in the CSF (LCC) and urine (Columbia) of PD patients compared to controls. **F.** Osteomodulin (OMD) protein intensity distribution in the urine of controls and PD patients of the Columbia cohort. Results of unpaired t test are shown. **G.** Correlation of ANCOVA q values of the proteins differentially present in the CSF (LCC) and urine (Columbia) of LRRK2 G2019S carriers compared to the LRRK2 WT controls. **H.** Correlation of ANCOVA q values of the proteins differentially present in the CSF (LCC) and urine (LCC) of LRRK2 G2019S carriers compared to the LRRK2 WT controls. **I.** CTSS protein intensity (log2) distribution in the urine of LRRK2 G2019S and WT carriers of the LCC cohort. Results of unpaired t test are shown.

Correlating the ANCOVA q values of all common proteins quantified in CSF (HBS and LCC) and urine (Columbia cohort) of PD patients compared to the controls revealed several proteins regulated in both fluids (**Figure 5D-E**). Among those was osteomodulin (OMD), the level of which was also significantly elevated in the urine of PD patients with a mean fold-change of 1.47 (**Figure 5F**). Next, we integrated ANCOVA q values of LRRK 2 G2019S vs LRRK2 WT of all common proteins quantified in the CSF samples of the LCC cohort and the urine samples of either Columbia or LCC cohorts (**Figure 5G-H**). This identified Cathepsin S to be regulated in a LRRK2 status-dependent manner in both matrices (**Figure 5G-H**). The levels of cathepsin S were significantly higher in the CSF of LRRK2 G2019S carriers compared to the LRRK2 WT carriers with mean fold-changes of 1.31 and 1.65 in the Columbia and LCC cohorts, respectively, (**Figures 5I and S2G**).

### Correlation of CSF proteome profiles with clinical scores indicating disease severity

We next investigated if any protein level changes in the HBS cohort correlated with the severity of PD pathology as assessed by the Unified Parkinson’s Disease Rating Scale (UPDRS) (**Figures 6 and S3, S4 and S5**). Indeed, 27 proteins are significantly correlated with the UPDRS scores in iPD patients (p-value < 0.001, **Figure 6A**). Proteins showing the highest positive correlation in PD patients included CHST6 (p value: 1.6E-5 and Pearson correlation r: 0.72), MIF (p value: 1.8E-5 and Pearson correlation r: 0.67), LYVE1 (p value: 4.8E5 and Pearson correlation r: 0.64), EFNA1 (p value: 5E5 and Pearson correlation r: 0.64) and ADM (p value: 1.3E4 and Pearson correlation r: 0.61) (**Figure 6A**). Proteins showing the highest negative correlation in PD patients included POMGNT1 (p value: 4.6E5 and Pearson correlation r: −0.64), TMEM132A (p value: 5.7E5 and Pearson correlation r: −0.63), ADAM22 (p value: 8.8E5 and Pearson correlation r: −0.62), PAM (p value: 1.9E4 and Pearson correlation r: - 0.6) and ST6GAL2 (p value: 2.7E5 and Pearson correlation r: −0.59) (**Figure 6A**). Interestingly, none of these proteins was significantly altered in a PD cases compared to controls. However, ADM was the strongest negatively correlated protein with the Mini-Mental State Examination (MMSE) scores (p value: 9.3E5 and Pearson correlation r: −0.43, **Figure S3D)**, a quantitative measure of cognitive impairment.

**Figure 6.**
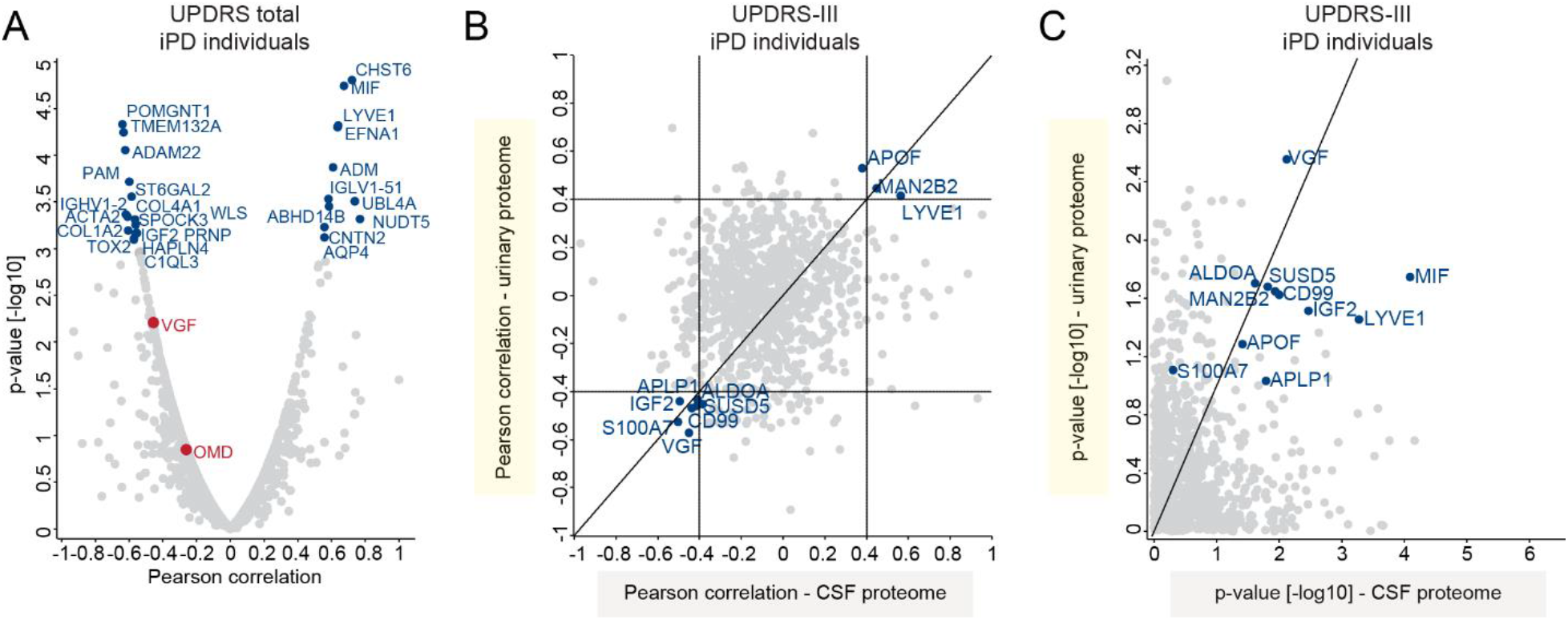
Correlation of CSF and urinary proteomes with UPDRS scores in iPD patients. **A.** Correlation analysis of protein intensities in CSF with the UPDRS total scores in iPD patients. Pearson correlation coefficients and -log10 p-values are displayed on the x- and y-axes, respectively. Proteins significantly correlating with UPDRS score (positively or negatively with a p-value < 0.001) are labeled. **B.** Correlation between Pearson correlation coefficients for correlation of urinary and CSF proteomes with UDPRS score **C.** Correlation between -log10 p-values for correlation of urinary and CSF proteomes with UDPRS score. The proteins with Pearson correlation coefficients > 0.4 in both datasets are labeled in B and C.

Furthermore, we compared the correlation scores of CSF proteins with those obtained in our previous study in urine. The comparison of Pearson coefficients for the UPDRS III scores (only UPDRS III scores were available for urine) revealed the 10 proteins APOF, MAN2B2, LYVE1, APLP1, ALDOA, IGF2, SUSD5, CD99, S100A7 and VGF to have coefficients either higher than 0.4 or lower than < −0.4 in both matrices (**Figure 6B**). Among these, VGF exhibited a significantly negative correlation with the UPDRS III scores in both datasets (p value of 2.8E3 and 7.6E3 in urine and CSF, respectively, **Figure 6C**). Overall, this analysis suggests that disease progression in PD patients, assessed by motor function, affects a similar set of proteins in CSF and urine.

Within the LCC cohort, Montreal Cognitive Assessment (MoCA) scores for all subjects were available (**Figure S1I)**. Interestingly, we found CHIT1 to significantly negatively correlate with MoCA scores in all patients (p value: 1.6E5 and Pearson correlation r: −0.48 in all subjects, **Figure S5D**) and this correlation was even stronger in PD patients (p value: 1.9E9 and Pearson correlation r: −0.81 in PD patients, **Figure S5E**). We also observed that two proteins strongly positively correlated with MoCA scores, RELN and PENK, were negatively correlated with age in CSF (**Figure S5D-E**). Thus, multiple proteins in the CSF proteome correlate well with disease severity and different clinically important aspects of the disease.

## DISCUSSION

Here we applied a scalable and highly reproducible MS-based proteomics workflow to CSF samples from two independent PD cohorts. To the best of our knowledge, our approach resulted in the deepest single-run CSF proteome acquired by mass spectrometry to date, with about 1,400 proteins identified and quantified per sample. In addition, our workflow is very sensitive, using less than 50 μl CSF for sample preparation. It does not require depletion of highly abundant proteins such as albumin or any biochemical enrichment, and the amount of purified peptides is sufficient for several MS runs, thus allowing the re-measurement of individual samples if required. Despite the unprecedented depth of our dataset, due to the high dynamic range of protein abundance in CSF, we have not yet identified some of the known and well-studied PD-relevant proteins such as α-synuclein and neurofilament light chain. Notably, the analytical variation of our assay, with a median CV of 19%, was much lower than the biological variation, with a CV of median 46%. We found this very precise, reproducible workflow well-suited to study disease-related biological differences.

To reduce systemic biases in the analyzed samples and minimize the effect of pre-analytical variation, we performed a thorough quality assessment of every sample using our previously reported quality marker panels [36]. We quantified the contamination of each CSF sample with platelets and erythrocytes and markers of potential coagulation events. This analysis flagged several samples of both analyzed cohorts as contaminated with erythrocyte-specific proteins and these were excluded from further analysis. Samples with a high degree of contamination were restricted to two of the four study centers. Erythrocyte counts, which are often determined following CSF sample collection, did not correlate with the degree of proteome contamination by erythrocyte-specific proteins, presumably because intact erythrocytes, which are typically determined in clinical labs, are frequently removed by centrifugation before samples are stored. Contaminations in the proteome are, however, not caused by intact erythrocytes but by hemolysis. Alternatively, we cannot exclude that different CSF aliquots were collected from each patient and that standard laboratory tests like erythrocyte counts were performed only on one of these aliquots. For the future, we recommend to collect collecting CSF in bulk first and aliquoting only following thorough mixing. Our findings using untargeted proteomics clearly underline the importance of stringent quality control and study protocols to avoid systemic biases, which may result in seemingly significant regulations in some quality markers that would then be reported as potential biomarker candidates.

ANCOVA analyses to identify proteins that are significantly regulated in PD patients compared to controls, identified CPM, OMD and RFNG in the HBS cohort and PRCP in the LCC cohort. The ANCOVA analysis considers confounding factors such as age, sex or LRRK2 status and thus is well suited to stringently assess which protein changes are truly associated with PD status. When we analyzed which proteins were disease status-dependently regulated in both cohorts, the OMD and CD44 stood out with very significant q values in both cohorts. OMD belongs to the small leucine-rich proteoglycans (SLRPs) and is involved in the organization and homeostasis of the extracellular matrix. CD44 is a cell surface glycoprotein involved in cell-adhesion and cell-cell interactions [46, 47]. Our results are in line with previous studies that identified OMD as a potential blood biomarker for PD [48] and upregulated in a SNCA transgenic mouse model [49]. A recent study has also shown the elevated expression of CD44 in the substantia nigra of human PD brains and CD44-mediated anti-inflammatory effects in primary mouse astrocytes [50]. Induced expression of CD44 by a-synuclein in microglia, likely affecting PD pathogenesis by recruiting reactive microglia into the pathological region of the PD brain, was also reported [51]. The low number of significantly regulated proteins in our study may be explained by the limited number of samples per disease group in each cohort combined with taking account of many potential confounders. Furthermore, patient heterogeneity due to inconsistent criteria for PD diagnosis and patient recruitment diminishes the overlap between independent cohorts. Future studies with larger sample cohorts are required to identify protein changes with small effect sizes.

A GO term enrichment analysis revealed that proteins upregulated in PD patients were associated with lysosome-related terms, which agrees with previous data that lysosomal dysregulation is evident in PD patients and involved in disease pathogenesis. In fact, lysosomal enzymes have long been investigated for their potential as biomarkers in the diagnosis of neurodegenerative disorders including PD [16, 52–56]. Changes in enzyme activities or abundances of lysosomal enzymes in CSF have been suggested to mirror the neuropathological changes linked to PD, although the basis of these alterations is not well understood. Levels of several lysosomal proteases including cathepsin D and cathepsin B are increased in CSF and post-mortem brain tissue of Alzheimer’s disease patients [57, 58]. Furthermore, activities of the lysosomal β-galactosidase and β-hexosaminidase as well as cathepsin L in CSF or postmortem brain tissue of PD patients are elevated [53, 55, 59–61]. Recently, we have shown that the pathogenic LRRK 2 - dependent changes of the urinary proteome included dozens of lysosomal proteins that could serve as biomarkers to stratify individuals with pathogenic LRRK2. In line with all these findings, in the present study, we identified two lysosomal proteins, Cathepsin K and MAN2B1, among the most important features to classify an individual’s PD status based on their CSF proteome. CTSS, another cathepsin, was one of the proteins with the highest upregulation in CSF of LRRK2 G2019S carriers compared to the LRRK2 WT controls in the LCC cohort. While cathepsin K has the potential to ameliorate α-syn pathology by degrading α-syn amyloids [62], increased expression of the CTSS gene – together with other genes involved in the antigen processing and presentation pathway and related immune pathways such as HLA-DQA1, HLA-DRA, HLA-DPA1, HLA-DMB - was reported in idiopathic PD patients in a study in which brain transcriptomic profiling was performed in idiopathic and LRRK2-associated PD [63]. Moreover, cathepsin S has been shown to regulate MHC class II antigen presentation process [64, 65].

Strikingly, several HLA proteins (HLA-DRA, HLA-DRB1, HLA-DPA) were significantly increased in LRRK2 G2019S carriers compared to controls. The HLA locus is one of the key loci associated with susceptibility for PD [15]. HLA-DR and HLA-DP are commonly expressed MHC-II molecules on antigen presenting cells, including microglia in the central nervous system [66]. Moreover, increased expression of HLA-DR is a hallmark of activated microglia, which are present in multiple neurodegenerative diseases including PD [66–69]. In addition, specific HLA-DRB1 variants can bind α-synuclein with high affinity and genome-wide association studies identified HLA-DRB1 and HLA-DRA alleles to be associated with PD in different populations [70–75]. A recent study has revealed a positive correlation between LRRK2 and MHC-II levels in PD patients and a negative correlation in healthy controls [76]. Our data support these findings and suggest a contribution of immunity and MHC-II molecules to the pathogenesis of familial PD.

Compared the analyzed CSF proteomes with our urinary proteome profiles in PD patients revealed 1080 common proteins. This large overlap and the clear correlation of the corresponding protein intensities (Pearson R 0.49) is remarkable and presumably due to shared origin of both CSF and urine, which is blood plasma. Interestingly, OMD was upregulated in PD patients in both CSF and urine, suggesting that the pathophysiology of PD affecting the OMD pathway is not restricted to the brain. CTSS was upregulated in LRRK2 G2019S carriers in both body fluids. LRRK2 is known to be ubiquitously expressed in many organs and the observed dysregulation of lysosomal enzymes including CTSS may be due to the hyperactivity of the mutated kinase. Moreover, we found VGF to be important in classifying PD patients from healthy controls by machine learning. Strikingly, we also reported this in our previous urinary proteome profiling study where VGF was the most important feature for classifying LRRK2+ PD patients from NMCs and its levels were strongly decreased in PD patients [27]. Consistently, VGF was also one of the handful of proteins found to be statistically significantly under-expressed in CSF of two independent PD cohorts [77]. Its negative correlation with the UPDRS-III scores of PD patients in both CSF and urine could indicate a protective role of this growth factor for motor function. Despite these interesting observations, the number of samples analyzed in our cohorts is still low for machine learning approaches and larger cohorts are needed to further improve the accuracy of the extracted models.

We also correlated clinical parameters with the CSF protein levels and identified several proteins including CHI3L1, FCGR3A, NCR3LG1 and ZP2 (p value < 0.01 in all analyses and both cohorts) that correlate well with the subjects’ age at sample collection (**Figures S3A-C and S5A-B**). CHI3L1 is a secreted glycoprotein that serves as a migration factor for astrocytes and a marker of glial inflammation [78, 79]. Interestingly, CHI3L was reported to be upregulated in the CSF of Alzheimer’s disease patients, therefore suggested as a biomarker for this disease and its expression levels in various regions of brain were shown to be correlated with age [26, 80–82]. Our data demonstrate that age correlates well with multiple proteins in the CSF and thus it is important to consider these confounders in statistical analyses to avoid a biased interpretation.

In line with our finding of CHIT1 being negatively correlated with the MoCA scores, chitinases including CHIT1 have emerged as biomarkers in neurological disorders as their levels correlate with disease activity and progression, likely reflecting microglia/macrophage activation [83, 84]. In addition, levels of proteins such as CHST6 correlate well with disease progression, as measured by the UPDRS (**Figures 6A and S4**). This correlation was particularly high for the UPDRS-III score, a measure of motor function. The sulfotransferase CHST6 plays an important role in keratan sulfonation and mutations in the corresponding gene cause macular corneal dystrophy [85]. Keratan sulfate, which is a type of sulfated Glycosaminoglycan (GAG), is also highly present in the brain, where it fulfils a multitude of functions [86]. CHST6 expression is increased in the brains of AD patients and its deficiency in mouse models mitigates AD pathology [87]. Interestingly, the sulfation state of GAGs affects α-synuclein aggregation by regulating lysosomal degradation [88]. Together, our data corroborate findings that sulfated GAGs like keratan sulfate can affect the pathophysiology of neurodegenerative diseases like PD. We identified a putative correlation between UPDRS-II scores and reduced LRPPRC expression, whose expression is also reduced in the blood of PD patients [89]. Mutations in *LRPPRC* cause the early-onset progressive mitochondrial neurodegenerative disorder French-Canadian-type Leigh syndrome, characterized by defects in oxidative phosphorylation reminiscent to those found in prodromal PD [90].

In conclusion, we have applied a highly reproducible and scalable MS-based proteomics workflow to perform proteome profiling of CSF in PD patients. We observed interesting and novel proteome changes in PD patients and identified biomarker signatures in the CSF that are specific to LRRK2 G2019S carriers. Our study provides further evidence that modern MS-based proteomics is a versatile and quantitative tool in biomarker discovery that can uncover pathological proteome changes in the CSF of patients suffering from a neurodegenerative disease like PD. Further studies analyzing larger cohorts of patients will be required to confirm our findings and extend the panels of potential biomarkers. In a next step, clinical and targeted assays then need to be developed to validate the biomarkers [12] and eventually develop a test that can be used in clinical routine to enable early disease detection and patient stratification.

## MATERIALS and METHODS

### Study cohorts

In this study, CSF samples from two independent cross-sectional cohorts were analyzed. Both studies were approved by local institutional review boards, and each participant signed an informed consent. The first cohort consisted of 94 CSF samples from the Harvard Biomarkers Study biobank and the second cohort was a subset of biobanked CSF samples from the Michael J. Fox Foundation for Parkinson’s Research (MJFF)-funded *LRRK2* Cohort Consortium (LCC). Disease severity was assessed by the Unified Parkinson’s Disease Rating Scale (UPDRS), which ranges from 0 for no impairment to a theoretical maximum of 199 for most severely affected individuals. UPDRS scores can be divided into four subscales for evaluating mentation, behavior, and mood (Part I, 0-16), activities of daily living (Part II, 0-52), motor examination (Part III, 0-108) and complications of therapy (Part IV, 0-23). Cognitive functioning was assessed using the Montreal Cognitive Assessment (MoCA), which ranges from 30 for no impairments to a theoretical minimum of 0 for most severely affected individuals. In addition, for the HBS cohort, Mini-Mental State Examination (MMSE) scores, which range from 30 to 0 (25-30 for normal cognition, 21-24 for mild dementia, 10-20 moderate dementia and 9 or lower for severe dementia) were available.

### Sample Preparation

40 μl of CSF samples were aliquoted in 96-well plates and processed with an automated set-up on an Agilent Bravo liquid handling platform as previously described [18, 26]. CSF samples were mixed with the equal amount of PreOmics lysis buffer (PreOmics GmbH) for reduction of disulfide bridges, cysteine alkylation, and protein denaturation at 95°C for 10 min. Upon 10 min cooling, 0.2 μg of each protease trypsin and LysC was added to each sample (well) and digestion was performed at 37°C overnight. Peptides were then purified on two 14-gauge StageTip plugs packed with styrenedivinylbenzene- reverse phase sulfonate (SDB-RPS) as described earlier [91]. The eluate was completely dried using a SpeedVac centrifuge at 45°C (Eppendorf, Concentrator plus), resuspended in 10 μl buffer A* (2% v/v acetonitrile, 0.2% v/v trifluoroacetic acid, and stored at −20°C. Upon thawing before mass spectrometric analysis, samples were shaken for 5 min at 2,000 rpm (thermomixer C, Eppendorf). Peptide concentrations were measured optically at 280nm (Nanodrop 2000, Thermo Scientific) and subsequently equalized using buffer A*. 500ng peptide was subjected to LC-MS/MS analysis.

Cohort-specific libraries were generated by pooling of 24 randomly selected samples of each cohort and separating the peptides of this pooled sample into 24 fractions each by high pH (pH 10) revers ed-phase chromatography as described earlier [92]. Fractions were concatenated automatically by shifting the collection tube every 120 seconds. Upon collection, fractions were dried and resuspended in buffer A* for LC-MS/MS analysis. To increase the depth of each library, we isolated extracellular vesicles (EV) from pooled CSF samples of each cohort by ultra-centrifugation as described earlier [93]. Isolated EVs were resuspended in 100 μl of a sodium deoxycholate-based lysis buffer containing chloroacetamide (PreOmics GmbH), heated to 95°C for 10 min for reduction and alkylation and then digestion using trypsin and LysC enzymes at 37°C overnight . Peptides were desalted with SDB-RPS StageTips as described earlier [91]. Peptides were eluted 80%/5% ACN/ammonium hydroxide and the eluate was completely dried and resuspended in 0.1% formic acid for separation into 8 fractions by high pH reversed-phase chromatography [92].

To determine coefficients of variation, five aliquots of a pooled CSF sample on one plate were subjected to sample preparation (intra-plate) and three aliquots of the same pool were subjected to sample preparation on two different plates (inter-plate).

### LC-MS/MS analysis

LC-MS/MS analysis was performed on a Q Exactive HF-X Orbitrap mass spectrometer with a nano-electrospray ion source coupled to an EASY-nLC 1,200 HPLC (all Thermo Fisher Scientific). Peptides were separated at 60 °C on 50 cm columns with an inner diameter of 75 μm packed in-house with ReproSil-Pur C18-AQ 1.9 μm resin (Dr.Maisch GmbH). Mobile phases A and B were 99.9/0.1% water/formic acid (v/v) and 80/20/0.1% acetonitrile/water/formic acid (v/v/v). MS data for single-shot CSF samples were acquired using the MaxQuant Live software and a data-independent acquisition (DIA) mode with phase-constrained spectrum deconvolution [94, 95]. Full MS scans were acquired in the range of m/z 300–1,650 at a resolution of 60,000 at m/z 200 and the automatic gain control (AGC) set to 3e6, followed by two BoxCar scans with 12 isolation windows each and a resolution of 60,000 at m/z 200 were acquired [96]. Full MS events were followed by 58 MS/MS windows per cycle in the range of m/z 300–1,650 at a resolution of 15,000 at m/z 200 and ions were accumulated to reach an AGC target value of 3e6 or for a maximum of 22 ms.

All fractionated samples including EV fractions were acquired using a top 12 data-dependent acquisition (DDA) mode. Full MS scans were acquired in the range of m/z 300–1,650 at a resolution of 60,000 at m/z 200. The automatic gain control (AGC) target was set to 3e6. MS/MS scans were acquired at a resolution of 15,000 at m/z 200.

### Mass spectrometry data processing

The MS data of the fractionated pools and the single-shot CSF samples were combined into two cohort-specific hybrid libraries using Spectronaut version 14.8.201029.47784 (Biognosys AG). For all experiments except the machine learning with OmicLearn, the two cohorts were quantified separately. All searches were performed against the human SwissProt reference proteome of canonical and isoform sequences with 42,431 entries downloaded in July 2019. Searches used carbamidomethylation as fixed modification and acetylation of the protein N-terminus and oxidation of methionines as variable modifications. The Trypsin/P proteolytic cleavage rule was used, permitting a maximum of 2 missed cleavages and a minimum peptide length of 7 amino acids. The Q-value thresholds for library generation and DIA analyses were both set to 0.01. For individual protein correlations with clinical parameters and the machine learning, the Q-value data filtering setting in Spectronaut was set to “Qvalue”. For all other analyses, the setting was set to “Qvalue percentile” with a cutoff of 25%, to use only those peptides for the protein quantification that passed the Q-value threshold in at least 25% of all analyzed samples.

### Data availability

The mass spectrometry proteomics data have been deposited to the ProteomeXchange Consortium via the PRIDE partner repository with the dataset identifier PXD02649, username: and password: 6rLsoARg.

### Bioinformatics data analysis

The Perseus software package versions 1.6.0.7 and 1.6.1.3 [97] and GraphPad Prism version 7.03 were used for the data analysis. Protein intensities were log2-transformed for further analysis apart from correlation and coefficient of variation analysis. Coefficients of variation (CVs) were calculated in Perseus for all inter-plate and intra-plate pairwise combinations of samples, the median values were reported as overall coefficient of variation. The protein CVs of the main study were calculated likewise within cohorts individually . For generation of the abundance curves, median protein abundances across all samples within a proteome were used. ANCOVA analysis was performed in python (version 3.7.6) using the pandas (version 1.0.1), numpy (version 1.18.1) and pingouin (version 0.3.4) packages. For the ANCOVA analysis, age at sample collection, *LRRK2* status (only in PD+ vs. PD-), and PD status (only LRRK2+ vs. LRRK2-) were set as confounding factors. The FDR was calculated using Benjamini-Hochberg correction. GO annotations were matched to the proteome data based on Uniprot identifiers. Annotation term enrichment was performed with Fisher exact test in Perseus separately for each cohort. Annotation terms were filtered for terms with an FDR of 5% after Benjamini-Hochberg correction in each cohort. Calculation of Pearson correlation scores and associated p-values of protein intensities to UPDRS scores and other clinical parameters was performed in Perseus.

### Machine learning

OmicLearn (v1.0.0) was utilized for performing the data analysis, model execution, and generating the plots and charts [40]. Spectronaut output tables from the quantification analysis of both cohorts were used as the input for OmicLearn. No additional normalization on the data was performed. To impute missing values, a Zero-imputation strategy was used. Features were selected using ExtraTrees (n_trees=500) strategy with the maximum number of 35 features. Normalization and feature selection was individually performed using the training data of each split. For classification, we used XGBoos t - Classifier (random_state = 23 learning_rate = 0.3 min_split_loss = 0 max_depth = 15 min_child_weight = 1). We used (RepeatedStratifiedKFold) a repeated (n_repeats=10), stratified cross-validat ion (n_splits=5) approach to classify PD vs. HC.

## ACKNOWLEDGEMENTS

Biospecimens used in the analyses presented in this article were obtained from the MJFF-sponsored *LRRK2* Cohort Consortium (LCC). For up-to-date information on the study: https://www.michaeljfox.org/news/lrrk2-cohort-consortium. We thank all members of the Proteomics and Signal Transduction Group at the Max Planck Institute of Biochemistry and the Clinical Proteomics Group at the NNF Center for Protein Research for help and discussions and in particular Jakob Bader, Philipp Geyer, Igor Paron, Johannes Mueller-Reif, Duc Tung Vu and Niels Skotte. We further thank Dario Alessi and Suzanne Pfeffer and their group members and employees of the Michael J. Fox Foundation for Parkinson’s research for helpful discussions.

The Harvard Biomarkers Study (“HBS”); https://www.bwhparkinsoncenter.org) is a collaborative initiative of Brigham and Women’s Hospital and Massachusetts General Hospital, co-directed by Dr. Clemens Scherzer and Dr. Bradley T. Hyman. The HBS Investigators have not participated in reviewing the current manuscript. The HBS Study Investigators are: Harvard Biomarkers Study. Co-Directors: Brigham and Women’s Hospital: Clemens R. Scherzer, Massachusetts General Hospital: Bradley T. Hyman; Investigators and Study Coordinators: Brigham and Women’s Hospital: *Yuliya Kuras, Karbi Choudhury, Michael T. Hayes, Aleksandar Videnovic, Nutan Sharma, Vikram Khurana, Claudio Melo De Gusmao, Reisa Sperling; Massachusetts General Hospital: John H. Growdon, Michael A. Schwarzschild, Albert Y. Hung, Alice W. Flaherty, Deborah Blacker, Anne-Marie Wills, Steven E. Arnold, Ann L. Hunt, Nicte I. Mejia, Anand Viswanathan, Stephen N. Gomperts, Mark W. Albers, Maria Allora-Palli, David Hsu, Alexandra Kimball, Scott McGinnis, John Becker, Randy Buckner, Thomas Byrne, Maura Copeland, Bradford Dickerson, Matthew Frosch, Theresa Gomez-Isla, Steven Greenberg, Julius Hedden, Elizabeth Hedley-Whyte, Keith Johnson, Raymond Kelleher, Aaron Koenig, Maria Marquis-Sayagues, Gad Marshall, Sergi Martinez-Ramirez, Donald McLaren, Olivia Okereke, Elena Ratti, Christopher William, Koene Van Dij, Shuko Takeda, Anat Stemmer-Rachaminov, Jessica Kloppenburg, Catherine Munro, Rachel Schmid, Sarah Wigman, Sara Wlodarcsyk; Data Coordination: Brigham and Women’s Hospital: Thomas Yi; Biobank Management Staff: Brigham and Women’s Hospital: Idil Tuncali.*

## AUTHOR CONTRIBUTIONS

OK and SVW designed the experiments, performed, analyzed and interpreted all data. DTV helped with analyzing the data. YK, IT, AMW and CRS were responsible for sample collection, their distribution and reviewed and edited the manuscript. SP, KM and CRS helped with interpretation of the results and reviewed and edited the manuscript. MM supervised the project and interpreted results. OK, SVW and MM wrote the manuscript.

## SUPPLEMENTARY FIGURES AND LEGENDS

**Figure S1.**
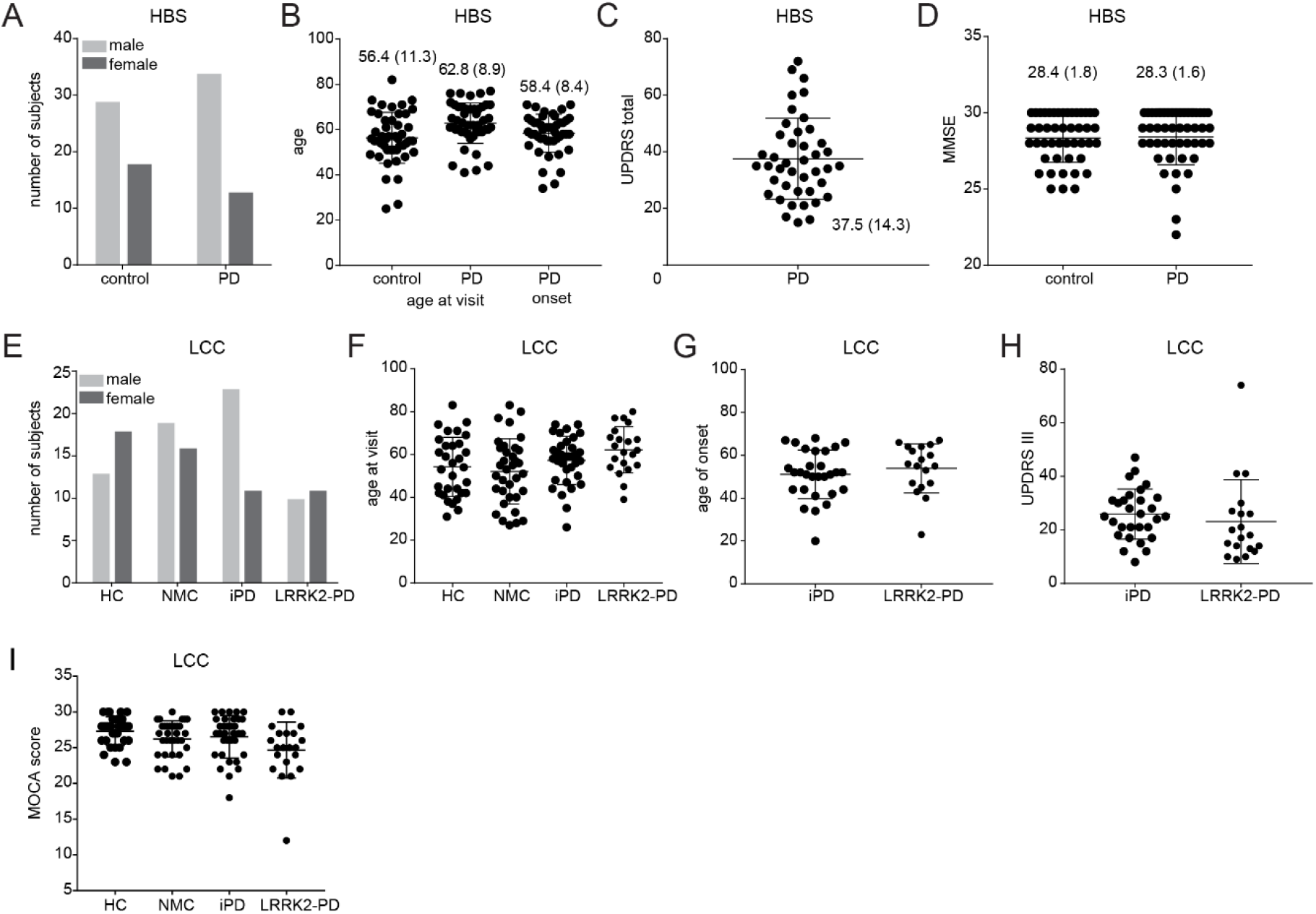
Clinical information and CSF proteome composition. **A**. Sex distribution among control and PD subjects in the HBS cohort. **B**. Age of subjects at the time of sample collection and disease onset for participants of the HBS cohort. Mean and standard deviation are shown. **C.** UPDRS total scores for PD individuals of the HBS cohort. Mean and standard deviation are shown. **D.** MMSE scores for healthy and PD individuals of the HBS cohort. Mean and standard deviation are shown. **E.** Distribution of subjects with different LRRK2 mutation status and manifestation of disease as well as sex in the LCC cohort. Number of subjects per group is shown. **F-G**. Age of subjects at the time of sample collection and disease onset for participants of the LCC cohort. Mean and standard deviation are shown. **H.** UPDRS III scores for iPD and LRRK2 PD individuals of the LCC cohort. Mean and standard deviation are shown. **I.** MOCA scores for all individuals of the LCC cohort. Mean and standard deviation are shown.

**Figure S2.**
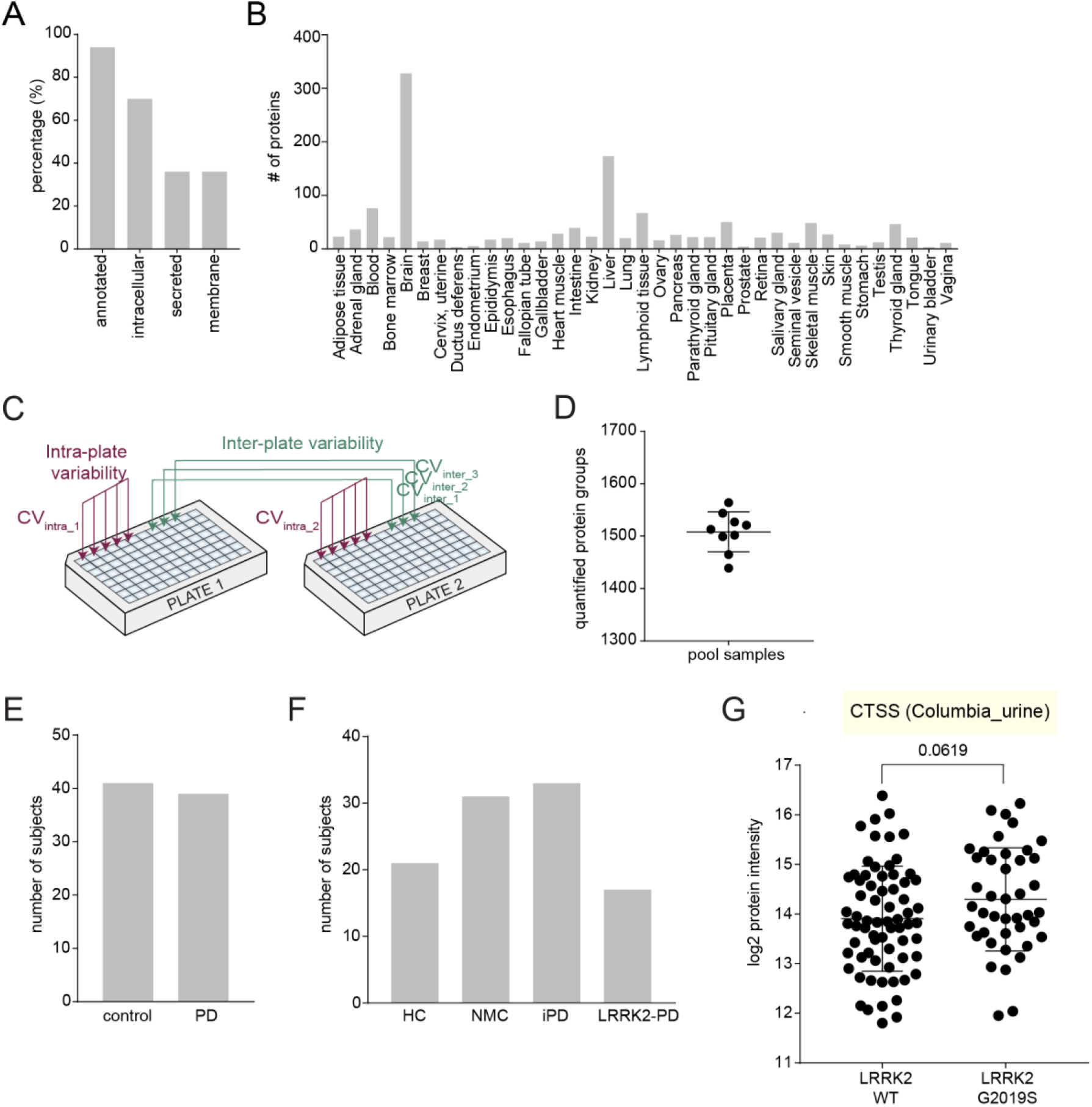
Assessment of the quantification precision. **A.** Percentage of identified CSF proteins which are secreted, located inside of the cell or in the cellular membranes. Proteins are annotated based on the Human Protein Atlas database. **B.** Number of tissue-specific proteins identified in CSF. The information regarding tissue specificities is retrieved from the Human Protein Atlas database (the Tissue Atlas). **C.** Graphical overview of the experiment to determine coefficients of variation (CVs) for the quantification of CSF proteins. **D.** Number of proteins quantified in each sample for the CV determination experiment. **E-F.** Distribution of samples after filtering the poor-quality ones according to erythrocyte contamination in the HBS (C) and LCC (D) cohorts. **G.** CTSS protein intensity (log2) distribution in the urine of LRRK2 G2019S and WT carriers of the Columbia cohort. Unpaired t test was applied and the resulting p values are shown.

**Figure S3.**
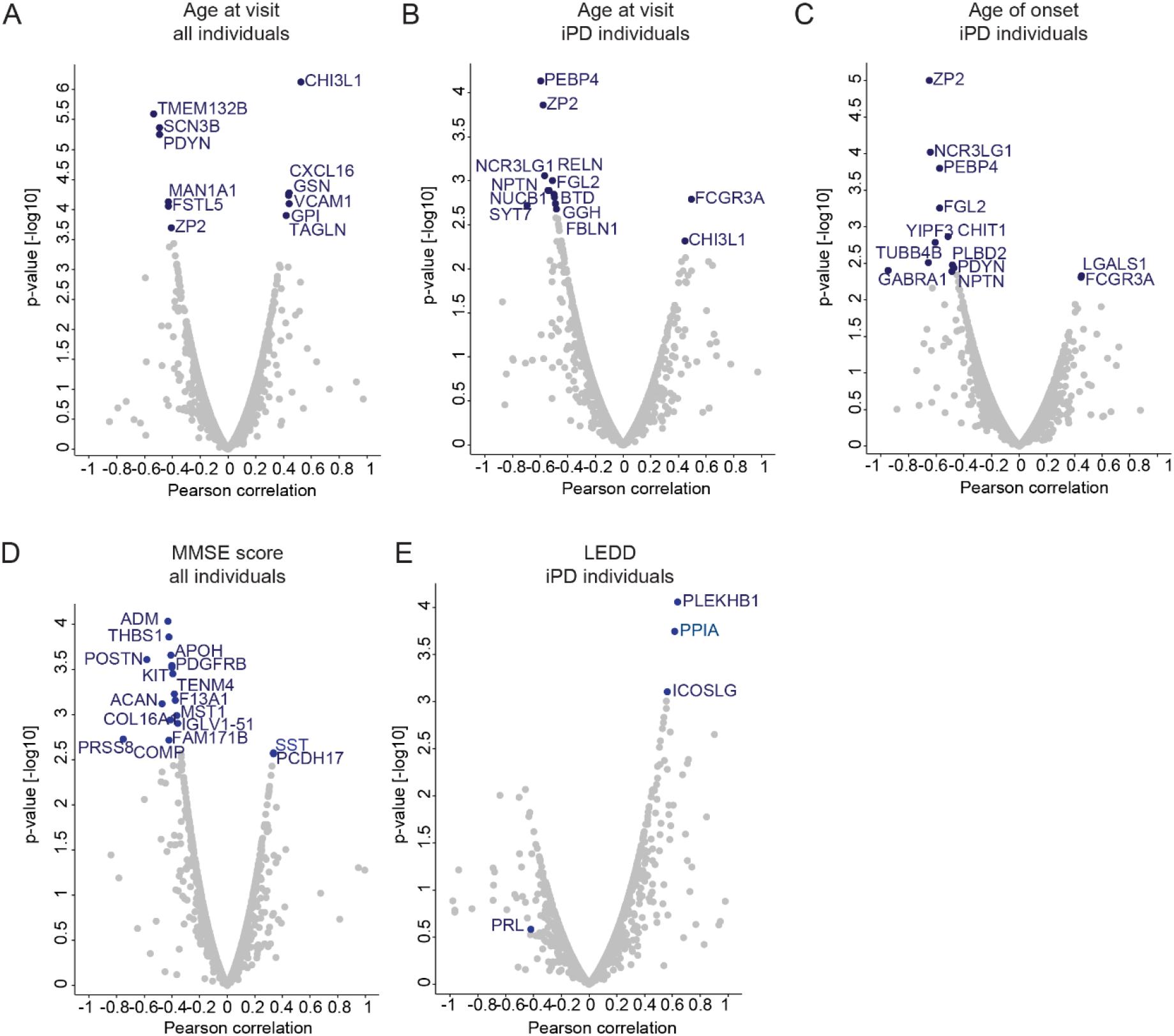
Correlation of protein abundances in the HBS with clinical parameters. **A-B**. Pearson correlation scores and associated p-values [-log10] of all protein intensities with the age at visit. All (A) or only iPD individuals (B) in the HBS cohort are included in the analyses as shown on top. **C.** Pearson correlation scores and associated p-values [-log10] of all protein intensities with the age of onset. All iPD individuals in the HBS cohort are included in the analysis. **D.** Pearson correlation scores and associated p-values [-log10] of all protein intensities with the MMSE score (**Figure S1D**). All individuals in the HBS cohort are included in the analysis. **E.** Pearson correlation scores and associated p-values [-log10] of all protein intensities with the LEDD levels. Only iPD cases in the HBS cohort are included in the analysis.

**Figure S4.**
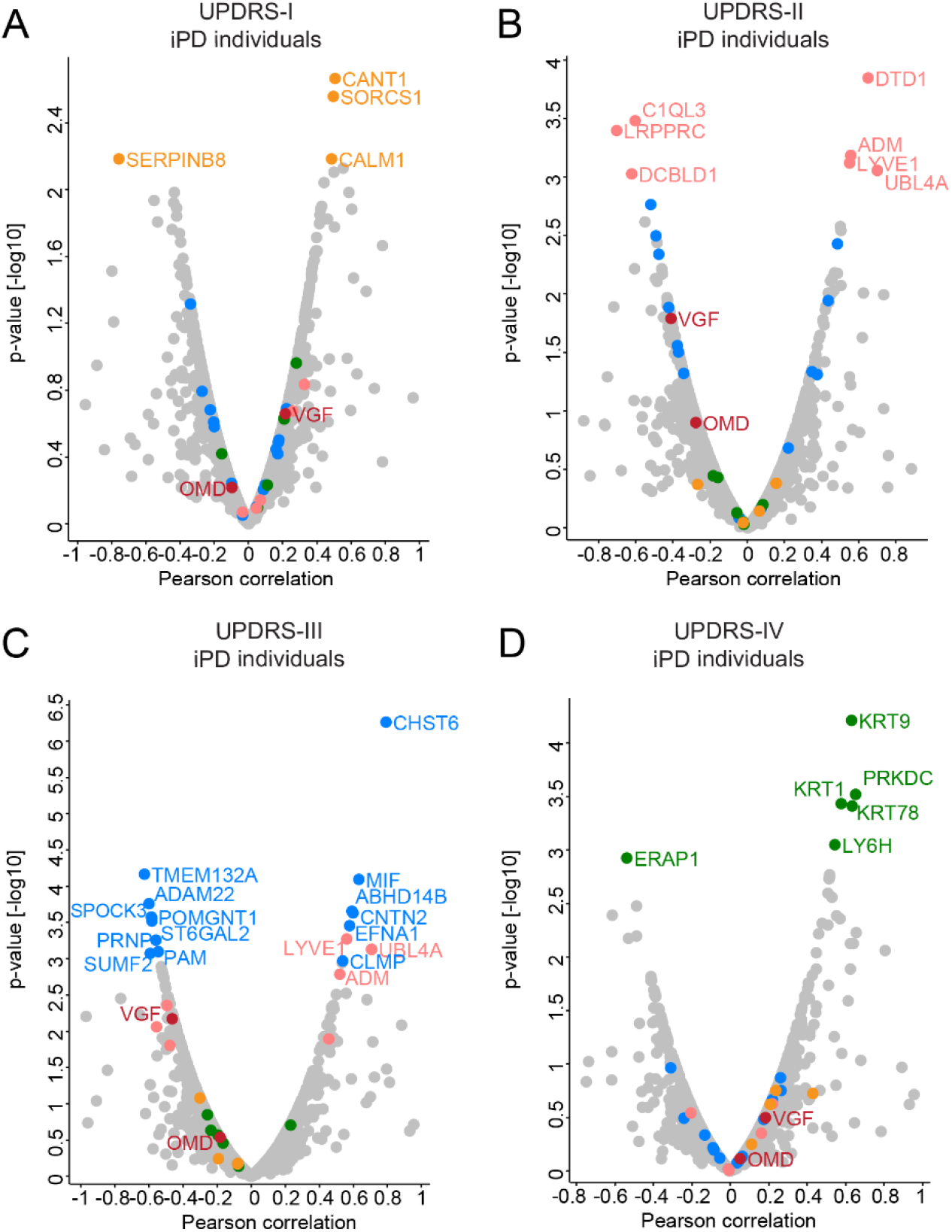
Correlation of protein abundances in the HBS with UPDRS scores. **A-D.** Correlation analysis of protein intensities in CSF with the UPDRS Part I (A), Part II (B), Part III (C) and Part IV (D) scores in iPD patients separately identified a different set of significantly correlated proteins. Pearson correlation coefficients and -log10 p-values are displayed on the x- and y-axes, respectively . Proteins significantly correlating with UPDRS score (positively or negatively with a p-value < 0.001) are labeled.

**Figure S5.**
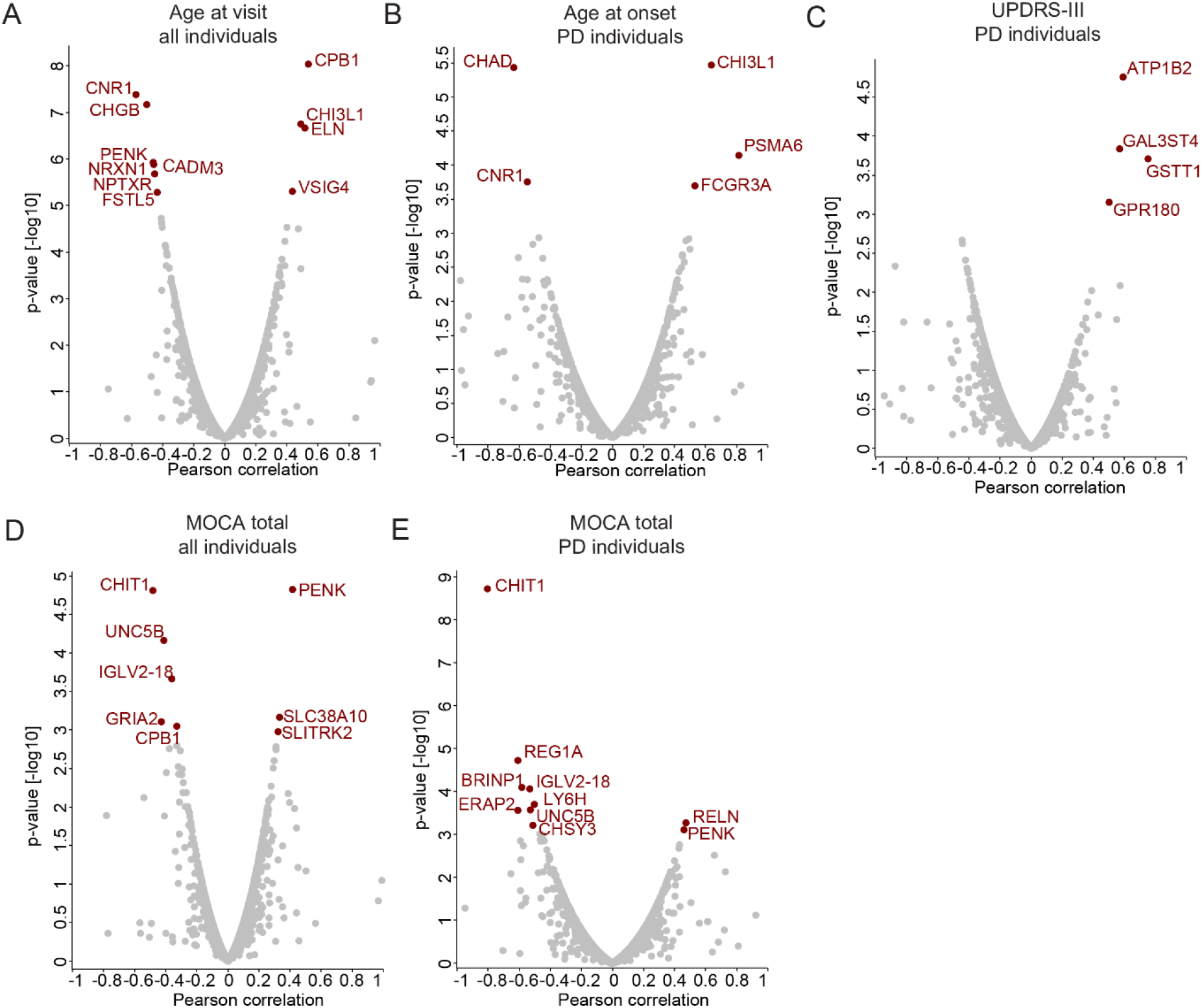
Correlation of protein abundances in the LCC with clinical parameters. **A**. Pearson correlation scores and associated p-values [-log10] of all protein intensities with the age at visit. All individuals in the LCC cohort are included in the analysis. **B**. Pearson correlation scores and associated p-values [-log10] of all protein intensities with the age of onset. PD patients in the LCC cohort are included in the analysis. **C.** Pearson correlation scores and associated p-values [-log10] of all protein intensities with the UPDRS-III scores (ranging 8 to 74) for PD patients included in the analysis. **D-E.** Pearson correlation scores and associated p-values [-log10] of all protein intensities with the MOCA scores. All individuals (D) or only PD patients (E) in the LCC cohort are included in the analyses as indicated on top.

**Figure S6.**
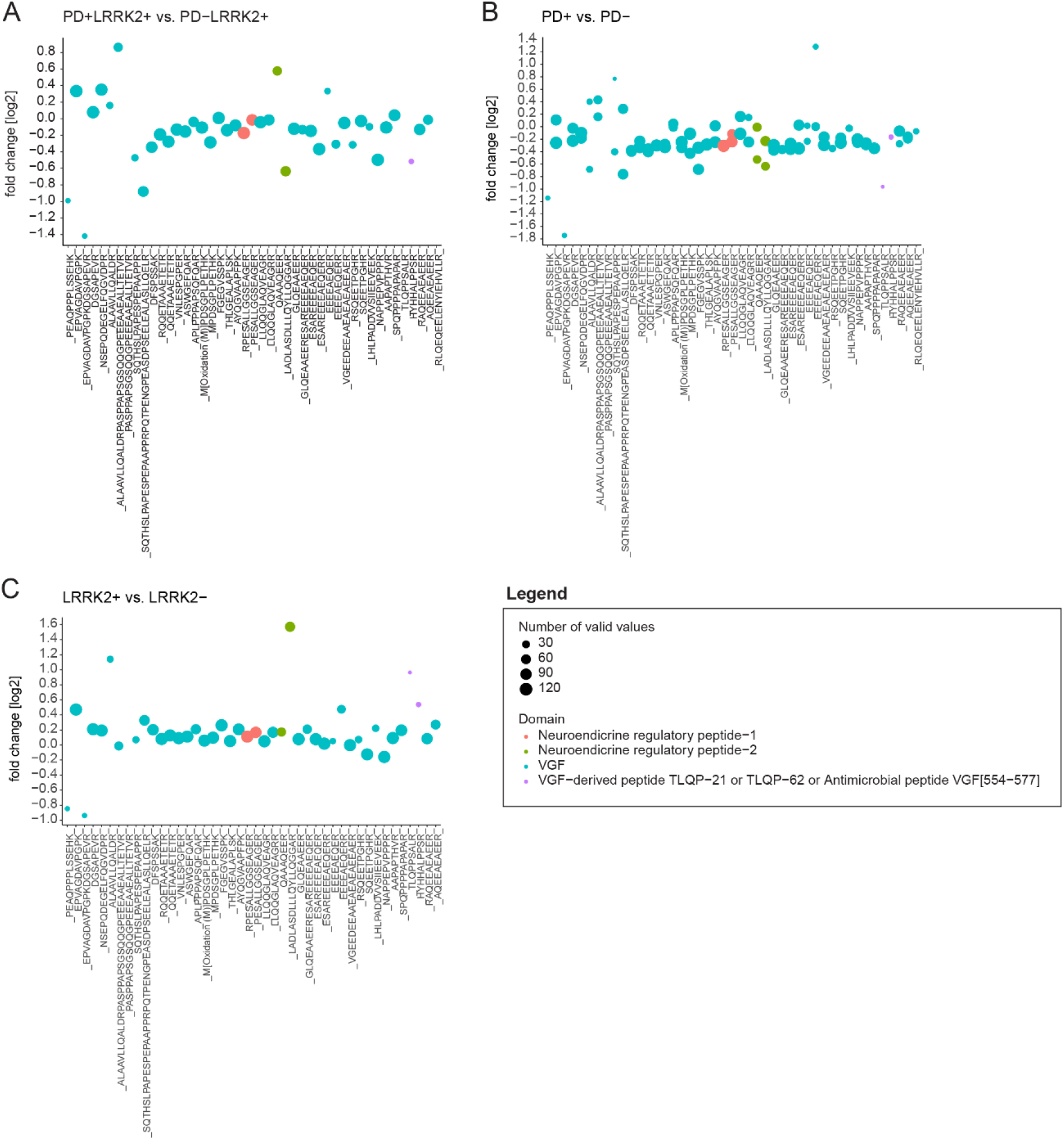
VGF peptides. **A**. Log2 fold-changes for each quantified VGF peptide comparing LRRK2+ PD patients with NMCs. The size of each dot represents the number of valid values (i.e. number of samples it was identified in). The color indicates if the quantified peptide sequence is part of a known functional VGF peptide. **B**. Log2 fold-changes for each quantified VGF peptide comparing PD patients and controls. **C**. Log2 fold-changes for each quantified VGF peptide comparing LRRK2 G2019S carriers and WT allele carriers.

